# Determine Protein Interaction Affinity without Purification Reveals a Two-enzyme Co-recognition of SUMOylation Substrate using Quantitative FRET (qFRET) Technology

**DOI:** 10.1101/2025.07.29.667540

**Authors:** Ling Jiang, Zhehao Xiong, Yan Liu, Runrui Dang, Jun Li, Jiayu Liao

## Abstract

Protein-protein interaction is the most fundamental process in life, and the quantification and manipulation of protein interaction activation or inhibition are critical for a comprehensive understanding of physiological processes and the development of new medicines that modulate protein-protein interactions. Due to its essential roles in physiology and pathology, many methodologies and technologies have been developed to determine the interaction events and parameters of interaction affinity. However, due to the nature of these methodologies, interactive partner needs to be purified, and large quantities of proteins are needed. This significantly limits quantitatively understanding of protein interaction and its application in therapeutics development. Recently, a quantitative FRET(qFRET) method was developed to determine purified protein-protein interaction affinity for purified proteins in vitro. Here, for the first time, we report the development of a high-throughput approach to determining the protein interaction affinity between two un-purified proteins using a quantitative FRET assay(qFRET). Two proteins, SUMO E2 conjugating enzyme, Ubc9, and RanGap1c, were first employed for the methodology development. We then determined the interaction affinities of Ubc9 and PIAS1 with influenza A virus NS1 protein without purifying two interactive partners. We found that both enzymes have high affinities to interaction affinity to the NS1, and this provides a kinetics explanation for the SUMOylation two-enzyme co-recognition mechanism to achieve high substrate specificity in vivo. The protein interaction affinity determination without purification provides several advantages over the current methods, including high accuracy, no tedious protein purification process, and high-throughput assay. The novel approach, so-called High-throughput determination of protein interaction affinity, KD, without purification using quantitative FRET assay, and scientific discoveries, representing a significant advancement in technology and scientific discovery, holds great potential for large-scale protein-protein interaction affinity determinations in vitro and in vivo.

**Teaser:** A novel qFRET technology for protein interaction affinity determination without protein purification for two-enzyme recognition.

## INTRODUCTION

Protein-protein interaction is central to all physiological processes, and alternation often occurs in many pathological developments, and therefore the determination and quantification of protein-protein interaction affinity are critical for understand living processes(*1, 2*). Due to their significant roles in life, many technologies, including Surface plasmon resonance(SPR), Calorimetric methods, such as ITC-isothermal titration calorimetry and DSC-differential scanning calorimetry, fluorescence polarization (FP), ultracentrifugation, radioactive labeling binding assay, were developed to determine protein interaction affinities(*3–5*). These methods have greatly stimulated investigations of protein functions and activities, ranging from ligand-receptor interaction, signal transduction, and enzyme-substrate recognition. However, these methods usually are time-consuming, labor-extensive and/or require expensive instruments, and low throughput measurements. Most significantly, current approaches require purified proteins and a relatively large quantity of proteins, which are often very challenging for many proteins and this significanly limits our understandings of protein interactions *in vitro* and *in vivo*. On the other hand, although several approaches, including tandem affinity purification with mass spectrometry (TAP-MS) and yeast/mammalian two-hybrid (Y2H) method, were developed to determine protein complex in the cells, protein interaction affinities in large-scale proteomics network and in cells are still not available, and the consequent quantitatively functionalities of most proteins in physiology and pathology remain poorly understood(*4, 6–9*).

Förster resonance energy transfer (FRET) is a nonradiative energy transfer that is based on the dipole–dipole interactions when the donor fluorophore (D) and acceptor fluorophore (A) have overlapped emission and excitation spectra and are close to each other (1–10 nm in general)(*10*). This distance is commensurate to the range of all biomolecular interactions in native status and can provide intra- and inter-molecular interaction information *in vitro* and *in vivo*. FRET has been widely used in biological/biomedical and translational research, including cell biology, biochemistry, diagnostics, optical imaging(*11–14*). FRET occurs. Due to the inverse sixth power dependence of FRET efficiency, FRET is very sensitive for nanoscale proximity detection with a typical range of 1-10nm(*15*). Therefore, qualitative and quantitative FRET-based imaging methods were developed to determine protein interaction in cells(*14, 16*).

The further developments of FRET assays into quantitative approaches in imaging and biochemical assays, mainly use quantitative three-cube or titration ratio-metric FRET assay to obtain FRET signals corresponding to the protein interaction events, which is correlated with protein interaction affinity, *K_D_*(*17*). However, the quantitative three-cube approach must determine the molar extinction coefficients of two fluorophores and many instrument-dependent parameters and estimate FRET efficiency during measurements, which often induces significant variations and makes it very difficult to turn the approach into an accurate one(*18, 19*). Another computational algorithm measured the intensity of individual voxels of 3-D FRET microscopy images in cells to infer the values of *K_D_*(*20*). However, this approach still needs calibration parameters to subtract background noise and does not consider endogenous protein interference. An *in vitro* FRET approach for *K_D_*determination used a calibration curve of point-to-point subtraction to obtain the fluorescence signals and ratio of acceptor and donor emissions as FRET signal, which did not exclude signals of direct emission of donor and acceptor during FRET assay(*21*). This ratio-metric FRET approach usually produces *K_D_* values larger than those determined by the SPR or the ITC. Multiple fluorescence parameter estimations and the difficulty of absolute FRET signal determination in above approaches limit their accuracy and robustness for *K_D_* determinations.

A recent development of quantitative FRET (namely qFRET) approach obtains the absolute FRET signal by subtracting the fluorescent emissions of the free donor and free acceptor from the total fluorescence emission in both purified protein assay (*22–25*). The emissions of free donor and acceptor are obtained from the donor or acceptor emissions multiplied by their cross-wavelength correlation constants, respectively. Then, the absolute FRET signal was used to correlate the protein-protein interactive complex at different acceptor concentrations to extrapolate the *K_D_* value in a two-parameter regression process. The *K_D_* value determined using this approach was in excellent agreement with the values determined by the SPR and the ITC approaches, demonstrating its accuracy and reliability. Furthermore, the qFRET assay is carried out in the 384-well plate, and therefore, it is the high-throughput assay mode. However, its potential for determining protein interaction affinity without purification, which represents a significant advancement, particularly for those difficult-to-be-expressed and purified proteins, has not been reported.

SUMOylation often changes protein activity, subcellular localization, and elongation of protein half-life and plays critical roles in transcriptional regulation, cell cycle, immune response, genome stability, epigenetics, neurodegeneration, and host-virus interactions(*26–29*). However, unlike Ubiquitination, SUMOylation E3 ligase is not absolutely required for SUMO conjugation for many substrates, especially in an *in vitro* SUMOylation reaction, but enhances SUMOylation and substrate specificity *in vivo*(*30, 31*). Generally, it is suggested that E3 ligase facilitates SUMOylation by orientating and stabilizing the Ubc9-SUMO complex to facilitate the transfer of SUMO peptide to the substrates(*31*). Out of the three families of E3 ligase for

SUMOylation, only the SP-Ring E3 ligase family, but not the RanBP2/Num358 and ZNF451 families, interacts with substrates directly(*31, 32*). The interaction of E3 ligase and substrates is often conducted via immunoprecipitation using E3-specific antibodies and immunoblotting with substrate-specific antibodies(*33, 34*). The SUMO E3 ligase-substrate interaction affinity, which can provide a mechanistic understanding of SUMO substrate recognition and contributions to substrate specificity, has never been determined.

Here, we report a novel development of qFRET analysis method to determine the protein interaction affinities of purified proteins without both protein purification in a high-throughput setting. For the first time, we also determine the protein interaction affinities of SUMOylation E3 ligase, PIAS1, and E2 conjugating enzyme, UBC9, to a SUMOylation substrate, influenza virus protein NS1. The discovery provides a novel kinetics basis for high specific recognition of substrate in SUMOylation, which can be invaluable for future anti-virus therapeutics developments. The innovative development of protein-protein interaction affinity determination without purification will enable us to understand protein interactions quantitatively in large-scale and in living organism in the future.

## MATERIALS AND METHODS

### DNA constructs

The CyPet and YPet sequences were cloned into pCRII-TOPO vector (Invitrogen) using the PCR with primers containing Nhe I-Sal I sites, respectively. RanGAP1c, Ubc9, NS1 and PIAS1 were amplified using PCR with primers containing Sal I and Not I sites. After sequencing conformation, the cDNA fragments encoding these genes were cloned into the Nhe I-Not I sites of pET28(b) vector for His-tag affinity purification (Novagen).

### Protein purification and concentration determination

The *Escherichia coli* BL_21_(DE3) strain was used for all protein expressions. After transformation, single colonies were picked up for seeding culture. Ten ml of cultures were added into 1L of 2xYT medium until bacterial cells grew to an optical density of 0.5–0.8 at 600 nm before the isopropyl β-D-thiogalactoside(IPTG) was added at a final concentration of 0.2 mM for an overnight induction. On the second day, bacterial cells were collected by centrifugation at 6,000 rpm for 10 min and resuspended in 30 ml of Ni^+^ column binding buffer (500 mM NaCl, 20 mM Tris-HCl, pH 7.4, and 5 mM imidazole). Then, the resuspended cells were sonicated for 5 min with 15-second sonication and 5-second intervals using an ultrasonic liquid processor (Misonix). The lysed cell lysates were then centrifuged at 35,000 g for 30 min to remove the insoluble fractions. The supernatants containing polyhistidine-tagged recombinant proteins were added to a 10 ml-column containing 1m of Ni^2+^-NTA agarose beads (QIAGEN), followed by three washes with the Washing buffer 1: 300 mM NaCl, 20 mM Tris-HCl, pH 7.4; Washing buffer 2: 1.5 M NaCl, 20 mM Tris-HCl, pH 7.4, and 5% Triton X-100; Washing buffer 3: 500 mM NaCl, 20 mM Tris-HCl, pH 7.4, and 20 mM imidazole). The bounded proteins were eluted by the Elution buffer (200 mM NaCl, 20 mM Tris-HCl, pH 7.4, and 250 mM imidazole). Finally, the eluted proteins were dialyzed against 1L of the Dialysis buffer (50 mM NaCl, 20 mM Tris-HCl, pH 7.4, and 1 mM DTT). The purity or protein mixtures were determined by the SDS-PAGE gel followed with Coomassie blue staining. The protein concentrations were determined using the Coomassie Plus Protein Assay Kit (Thermo Scientific).

### Fluorescence signal determination in 384-well plate

All the FRET assays were performed in 384-well plates and the fluorescence was determined using the fluorescence plate reader FlexstationII384 (Molecular Devices, Sunnyvale, CA). Specifically, to determine the correlation coefficient constant, α, and total fluorescence emission signal, Em_Total_, the fluorescence emission signals at 475 and 530 nm, respectively, were determined when the excited at 414 nm with a cutoff filter at 455 nm. To determine the correlation coefficient constant β, YPet-conjugated proteins’ fluorescence emission at 530 nm was determined when excited at 495 nm with a cutoff filter at 515 nm. All the assays were performed in triplicate, and the average fluorescence signal value was recorded under each condition for standard deviation calculation.

We performed FRET assays in four different settings: purified proteins, purified proteins plus BSA, purified proteins in the presence of bacterial cell extracts, and unpurified fluorescent protein mixtures. In all the settings, in order to make sure the assays could give consistent results, several concentrations of CyPet-proteins were set up, in general, four concentrations as 0.05, 0.1, 0.5, 1.0 µM, and ten concentrations of the YPet-proteins from 0 to 4 µM were added to research plateaus of the titration curve. In the first FRET assay, purified CyPetRanGAP1c and YPetUbc9 proteins were mixed in the Interaction buffer (150 mM NaCl, 20 mM Tris-HCl, pH 7.5, DTT 1 mM) in a total volume of 60 µL. In the second FRET assay, one ug of pure BSA was added to the purified CyPetRanGAP1c and YPetUbc9 protein mixture in the Interaction buffer. In the third FRET assay, purified CyPetRanGAP1c and YPetUbc9 proteins were mixed with a bacterial extract made from BL_21_(DE3) E.coli cell after sonication and centrifuge. In the fourth FRET assay, unpurified CyPetRanGAP1c, YPetUbc9, CyPetNS1, YPetNS1, and YPetPIAS1 proteins were obtained from BL_21_(DE3) E.coli cells expressing each individual protein after sonication and centrifuge without purification. We then measured the concentrations of unpurified proteins using the Fluorescence standard curves of CyPet and YPet, respectively(see below). In the FRET assay, the final concentrations of CyPetRanGAP1c were fixed at 0.05, 0.1, 0.5, and 1.0 µM, respectively, and the final concentrations of YPetUbc9 were varied from 0 to 4 µM.

### Establishment of the standard curves of CyPet-protein and YPet-protein

To establish external fluorescence emission standard curves, the different concentrations of the purified recombinant protein CyPet or YPet in the Interaction buffer were added to the 384-well plate in a total volume of 60 µL. The concentration of CyPet protein varied from 0 to 2 µM, and the emission signals of 475 nm were measured when excited at 414 nm. The concentration of YPetUbc9 protein ranged from 0 to 2 µM, and the emission signals at 530 nm were determined when excited at 475nm. BL21 bacteria cell lysate was used as a background control. The fluorescence emissions were measured in a 384-well black plate using FlexstationII384 (Molecular Devices, Sunnyvale, CA).

### FRET spectrum analysis for absolute FRET signal determination

The general procedure of fluorescence spectrum analysis for FRET assay was described before(*22, 24, 25*). Briefly, when the FRET pair-labeled interactive proteins were excited at 414 nm, peak emission at 475 nm defined as FL_DD_ and 530 nm defined as Em_total_ were measured (see Fig. 1A). The acceptor emission at its excitation wavelength defined as FL_AA_ (for YPet, emission signal at 530 nm when excited at 495 nm) was determined. In the FRET assay of CyPet-protein and YPet-protein mixture, when excited at 414 nm, almost all of the emission signal at 475 nm comes from the direct emission of CyPet-proteins because the YPet protein cannot be excited at 414nm. On the other hand, the emission signals at 530 nm (Em_total_) comes from three contributions: the direct emission of CyPet-protein, the direct emission of YPet-protein, and the sensitized absolute FRET signal of YPet-protein(Em_FRET_). Because Em_FRET_ is proportional to the amount of the formed interactive complex, we can develop mathematic algorithm between Em_FRET_ and the bound protein concentration to determine *K_D_*value(*24*).

**Figure 1.**
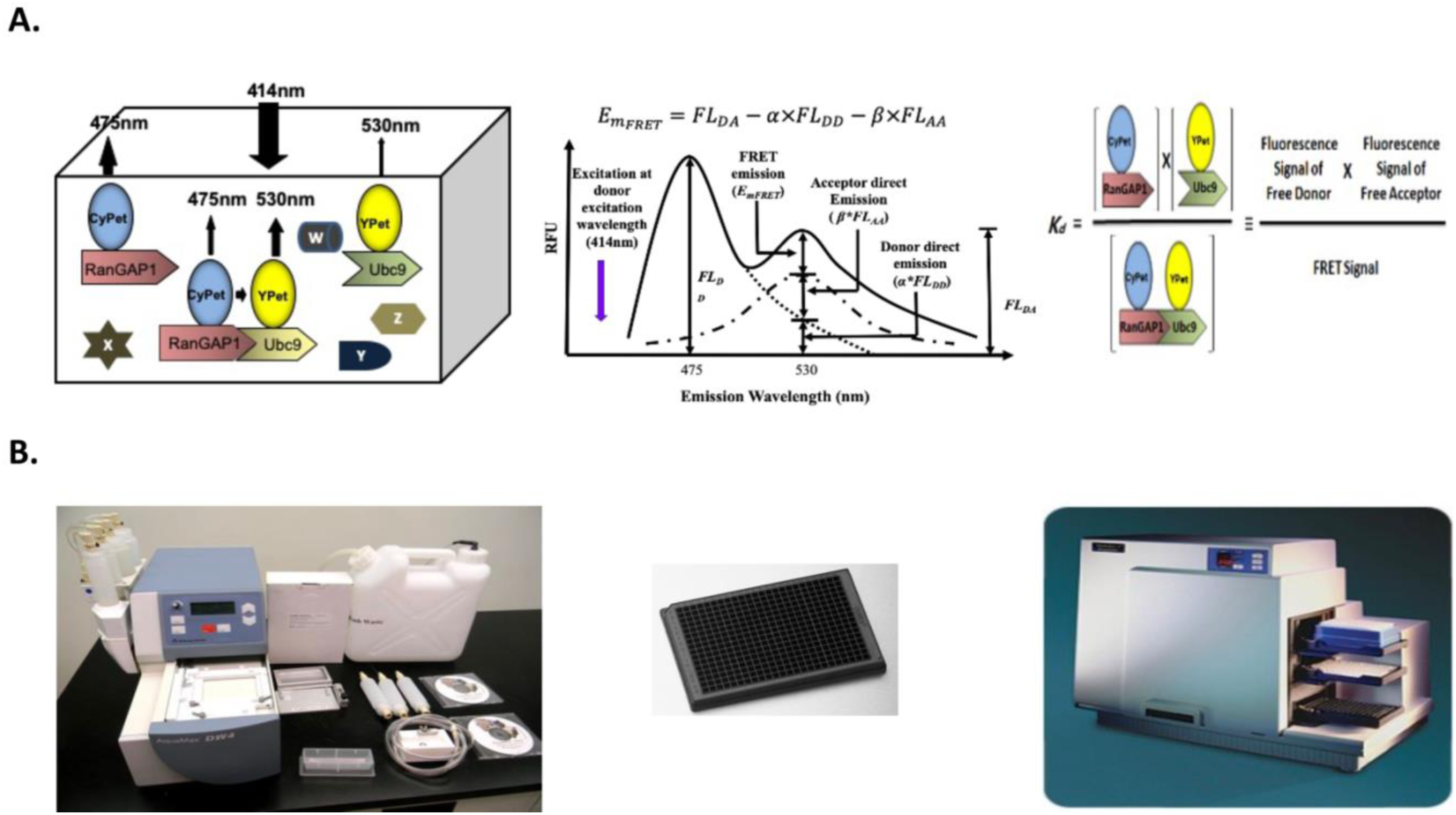
High-throughput FRET-based technology for protein interaction dissociation constant, *K_d_*, determination without purification. A. Schematic graph of fluorescence excitation and emission signals of interactive proteins, CyPet-RanGAPs and YPet-Ubc9, in the presence of other proteins(Left); quantitative FRET(qFRET) signal determination(Middle); and protein interaction dissociation constant, *K_D_*, determination based on fluorescent signals of free donor and acceptor as well as interaction complex. B. High-throughput FRET assay for semi-automatic fluorescent protein distribution(Left); 384-well FRET assay platform(Middle); and Fluorescence plate reader, Flexstation^384^.

During the experimental procedure to determine the sensitized absolute FRET signal, Em_FRET_, two experiments were first conducted to determine the cross-wavelength correlation coefficient ratio constants, *α* and *β*. The first experiment was conducted to determine the correlation coefficient constant of *α* for the donor. Different concentrations of CyPetRanGAP1c protein, 0.05, 0.1, 0.5, and 1.0 µM, were measured for theemissions at 475 and 530 nm when excited at 414 nm. Dividing the fluorescence emission of the donor CyPetRanGAP1 at 530 nm (FL_DA_) by its emission at 475 nm (FL_DD_) yielded the correlation coefficient ratio constant, *α*. The average α values at different concentrations are used as the final value α. The α is used to estimate fluorescence emission of the donor CyPetRanGAP1c at 530 nm when excited at 414 nm. In the second experiment, a series of YPetUbc9 protein concentrations, 0.2, 0.5, 1, 2, 3, and 4 µM, were measured for their emission at 530 when excited at 495 nm. The correlation coefficient ratio constant *β* is defined as the fluorescence emission at 530 nm when excited at 414 nm(FL_AD_) divided by its emission at 530 nm when excited at 495 nm (FL_AA_). The average β values at different concentrations are used as the final value β. The β is used to estimate the fluorescence emission of the acceptor YPet at 530 nm when excited at 414 nm. The final absolute FRET signal can then be determined using the following equation,

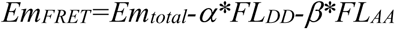

where the *Em_total_, FL_DD_, FL_AA_* were determined during the FRET assay, while the α and β were pre-determined.

### Data processing for *K_D_* determination

After the FRET signal (Em_FRET_) was determined as described above, the dataset of Em_FRET_ and the corresponding concentrations of [YPetUbc9]_total_ was fitted to the following equation to derive Em_FRETmax_ and *K_D_* through the multiple variable regression using Prism 5 (GraphPad Software),

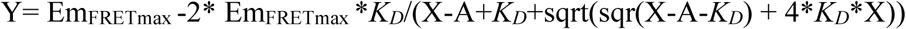

Specifically, the values of [YPetUbc9]_total_ were plotted into X-series and the Em_FRET_ intensities at each concentration of [YPetUbc9]_total_ were plotted into Y-series in the above equation. A nonlinear two-variable regression method was applied using the above equation to fit the dataset. The initial values for the parameters of Em_FRETmax_ and *K_D_* were set as 1.0, and the default constraint of the Em_FRETmax_ must be greater than 0. The *K_D_*and Em_FRETmax_ values were reported as mean ± SD.

### Determination of interaction affinity *K_D_* values using Surface Plasmon Resonance (SPR)

The YPetUbc9, CyPetRanGAP1c, Ubc9 and RanGAP1c proteins were expressed and purified from E.coli strain Bl_21_(DE_3_) and then dialyzed in the Binding buffer (150 mM NaCl, 10 mM HEPES, 50 µM EDTA, 0.005% Tween20, pH7.4). All SPR measurements between RanGAP1c and Ubc9 or CyPetRanGAP1c and YPetUbc9 were conducted using an NTA sensor chip at a flow rate of 30 µL/min using the BIAcore X100 system (BIAcore AB, Uppsala, Sweden). The chip was first treated with 500 µM of NiCl_2_ solution in the Binding buffer for 1 min. To immobilize proteins on the chip, 100 ng/mL of purified YPetUbc9 or 200 ng/mL of purified Ubc9 protein was injected into the chips. In the binding and dissociation phase, a ∼100 µg/mL of thrombin-digested CyPetRanGAP1c protein or ∼25 µg/mL of thrombin-digested RanGAP1c protein was injected into the chip for 120s and disassociated for 10min to each of corresponding interactive partner-immobilized chip. As a nonspecific control, To ensure a low background binding of samples to the NTA chip surface, CyPetRanGAP1c and RanGAP1c proteins were injected into a control flow cell without treatment of NiCl_2_ and YPetUbc9 or UBC9 proteins. After one round of experiments, the NTA sensor chip was regenerated using a buffer (350 mM EDTA, 150 mM NaCl, 10 mM HEPES, 0.005% Tween20, pH8.3), and then reloaded with NiCl_2_. Data were analyzed using BIAcore X100 software (Ver.1.0, BIAcore).

### Statistical Analysis

All the *K_D_* data under each donor concentration and assay condition combination are reported as the mean ± standard error (SEM). The two-variable design offers the advantages of higher experimental efficiency and the capability of statistically analyzing the interaction of the two factors. The two-way ANOVA test provides a statistical analysis to determine how the *two factors affect the KD value*s. Here, we implemented a two-way ANOVA analysis using GraphPad Prism to compare the *K_D_* values under different combinations of donor concentration (row factor) and assay conditions (column factor). This analysis divides the total variability among *K_D_* values into four components: interactions between row and column, variability among columns, variability among rows, and variability among replicates. It computes three P values, which test three null hypotheses, respectively:

□ The interaction P value to test the null hypothesis is that there is no interaction between columns and rows.

□ The column factor P value to test the null hypothesis is that the mean of each column is the same.

□ The row factor P value to test the null hypothesis is that the mean of each row is the same.

## RESULTS

### Theory of <u>h</u>igh-throughput determination of <u>p</u>rotein interaction <u>a</u>ffinity, *K_D_*, from <u>m</u>ixture proteins without purification using quantitative <u>F</u>RET assay (HPAMF)

We have previously developed a quantitative FRET(qFRET) method to elucidate the FRET signal responding to the protein interactive complex to determine the *K_D_* from either acceptor emission or donor quenching(*22, 35, 36*). But we never explored this method to determine protein interaction affinity in an environment with other proteins or even from crude cell extract without purification (Fig.1A, left) using the qFRET analysis to differentiate sensitized FRET fluorescence signal resulted from protein interactive complex from those of free ligand and free acceptor (Fig.1A, middle), following the Mass action principle (Fig.1A, right). This is not just a simple extension of the previous approach but advances this methodology to another level of protein interaction affinity determinations for difficult-to-be-expressed/purified proteins. Because other proteins do not contribute to the fluorescence signal as compared with the strong FRET pair proteins,

Briefly, in the method of determining interaction affinity *K_D_*, the critical step is to obtain the absolute sensitized FRET signal (E_mFRET_) corresponding to the interactive protein complex in order to determine the maximum bound protein complex regardless of purified or unpurified proteins. The total fluorescence emission at the FRET emission signal wavelength(E_mTotal_), in the case of 530nm for the FRET pair CyPet and YPet, consists of three components: the direct emission of the non-bound acceptor, YPet, which comes from the free acceptor in the mixture, the direct emission of the none-bound donor, CyPet, which comes from the free donor not in the interactive complex, and the absolute sensitized FRET signal (E_mFRET_), which comes from the protein interaction complex (Fig. 1A, middle). Therefore, if we can determine the direct emissions of the free donor and free acceptor, we can obtain the absolute sensitized FRET signal (E_mFRET_). The direct emissions of the donor and acceptor, named FLDD and FL_AD_, respectively, can be determined using a cross-wavelength correlation coefficient method.. The structure of the fluorophores in the fluorescence proteins determines the fluorescence emission spectrum, which is an intrinsic property of each fluorescence protein. Therefore, we can determine the direct emission of the donor CyPet at 530nm by timing its emission at 475nm when excited at 414 nm with aconstant ratio α, which is defined as *α* =FL_DA_/FL_DD_. Using a similar approach, we can determine the direct emission of the free acceptor YPet at 530nm by timing its emission at 530nm when excited at 495 nm with a ratio factor β, which is defined as *β* =FL_AD_/FL_AA_. The FL_AA_ is the fluorescence emission of Acceptor, YPet, at 530 nm when excited at the acceptor excitation wavelength, 495 nm) and FL_AD_ is the acceptor emission at acceptor emission wavelength when excited at the donor excitation wavelength. In the experimental procedure, first, when the mixture of interactive partners, CyPetRanGAP1c and YPetUbc9, is excited at the donor excitation wavelength (414nm in this case), two emission signals at 475nm (FL_DD_) and 530nm (FL_DA_) are determined. The fluorescence emissions at 475nm when excited at 414 nm come from both the emission of CyPet (FL_DD_) and the direct emission of YPet at 475 nm. Then, the FRETemissionsignalof the interactive complex, CyPetRanGAP1c/YPetUbc9 in this study, can be determined by subtracting the above two signals, ***α*\***FL_DD_ and β*FL_AA_, from the total emission at 530 nm, E_mTotal_, or the FRET emission signal (Em_FRET_) can be determined using the following formula:

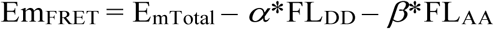

where the ratio constants *α* and *β* are first experimentally determined as 0.334±0.003 and 0.014±0.002, using free CyPetRanGAP1c and YPetUbc9 in this case, respectively.

After the Em_FRET,_ is obtained, following the general Mass action law,

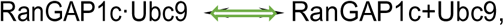

The protein interaction dissociation constant, *K_D_*, can be determined as the following ,

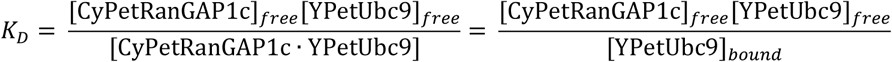

This can be rearranged to

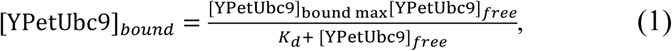

where [YPetUbc9]_bound max_ is the theoretical maximal bound complex of YPetUbc9 and CyPetRanGAP1c when donor CyPetRanGAP1c concentation is fixed in each assay, and [YPetUbc9]_free_ is free YPetUbc9 concentration. [YPetUbc9]_bound_ is proportional to the FRET signal from the bound protein complex. Then Eq.(1) can be converted into Eq.(2) using the relationship,

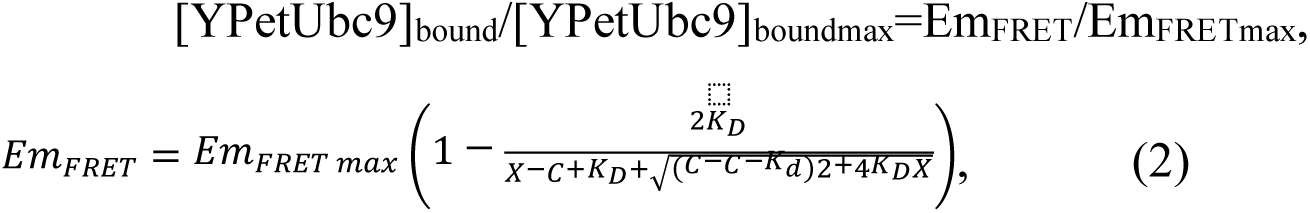

where Em_FRET_ is the absolute FRET signal, and Em_FRET max_ is the theoretical absolute FRET signal when YPetUbc9 binds the maximum amount of CyPetRanGAP1c in the assay. The C is the total concentration of CyPetRanGAP1c in the assay, and X is the concentration of total YPetUbc9 added in each of the assay. Using this equation, a multiple-variable regression approach can determine the K_D_ and Em_FRET max_.

In the study, all the fluorescence measurements were performed in a 384-well plate using the fluorescence plate reader FlexStaion384 (Fig.1 B). Therefore, this assay is conducted in a high-throughput assay format.

### Determine interaction affinity *K_D_* of purified Ubc9 and RanGAP1c using HPAMF

We first determined the E_mFRET_ and *K_D_* of purified Ubc9 and RanGAP1c to establish the interaction’s reference parameters for a later methodology development. One concentration of donor is sufficient to determine the interactive dissociation constant *K_D_*. But to test the generality of the method, the CyPetRanGAP1c was set up as 0.05 µM, 0.1 µM, 0.5 µM and 1.0 µM four concentrations, and in each assay, the acceptor YPetUbc9 concentration was titrated from 0 to 4 µM. The interaction of YPetUbc9 and CyPetRanGAP1c led to an energy transfer from CyPet to YPet, in which the fluorescence emission at 530 nm was increased, so-called FRET emission, while the emission of the donor CyPetRanGAP1c at 475 nm decreased, so-called FRET quenching. We then determined the fluorescence signals, E_MTotal_, FL_DD_, and FL_AA_, in four conditions of CyPetRanGAP1c. The α and β were pre-determined using purified proteins CyPetRanGAP1c and YPet-Ubc89, respectively. After subtracting the direct emissions of CyPetRanGAP1c (***α*\***FL_DD_) and YPetUbc9 (***β***\*FL_AA_) at 530 nm from the total emission at 530 nm (E_MTotal_), the absolute FRET signals at 530 nm when excited at 414 nm (Em_FRET_) were determined as an increase steadily when more YPetUbc9 was added at each concentration of CyPetRanGAP1c. The values for Em_FRETmax,_ were calculated as (1.23±0.02)x10^4^, (2.43±0.04)x10^4^, (12.29±0.23)x10^4^, and (24.61±0.53)x10^4^, for 0.05, 0.1, 0.5, 1.0 µM of CyPetRanGAP1c, respectively(Supplement Fig.1 A) (Supplement Table 1). In this concentration range of the binding partner, the Em_FRETmax_ had a linear relationship with CyPetRanGAP1c from 3 to 60 pmole (Supplement Fig. 1B). This result suggests that our approach accurately and consistently predicted Em_FRETmax_ at various concentration ratios of CyPetRanGAP1c and YPetUbc9.

The disassociation constants of CyPetRanGAP1c and YPetUbc9 were then determined from the non-linear regression. By plotting Eq.(2) with Em_FRET_ vs. [YPetUbc9]_total_ in the Prism 5 program, we determined *K_D_*s from four concentrations of CyPetRanGAP1c (0.05, 0.1, 0.5, and 1.0 μM) as 0.098±0.014, 0.096±0.013, 0.101±0.016, and 0.114±0.021 μM, respectively, which were very close to the *K_D_* value determined using SPR, , 0.097μM, (Supplement Table 1) (Table 1). The very close values of *K_D_*generated from different concentrations of CyPet-RanGAP1c(from 0.05 μM to 1.0 μM of CyPetRanGAP1c) and various binding partner ratios of CyPetRanGAP1c to YPetUbc9 (from 0.67 to 40 folds) demonstrate that this method is very robust.

**Table 1.**
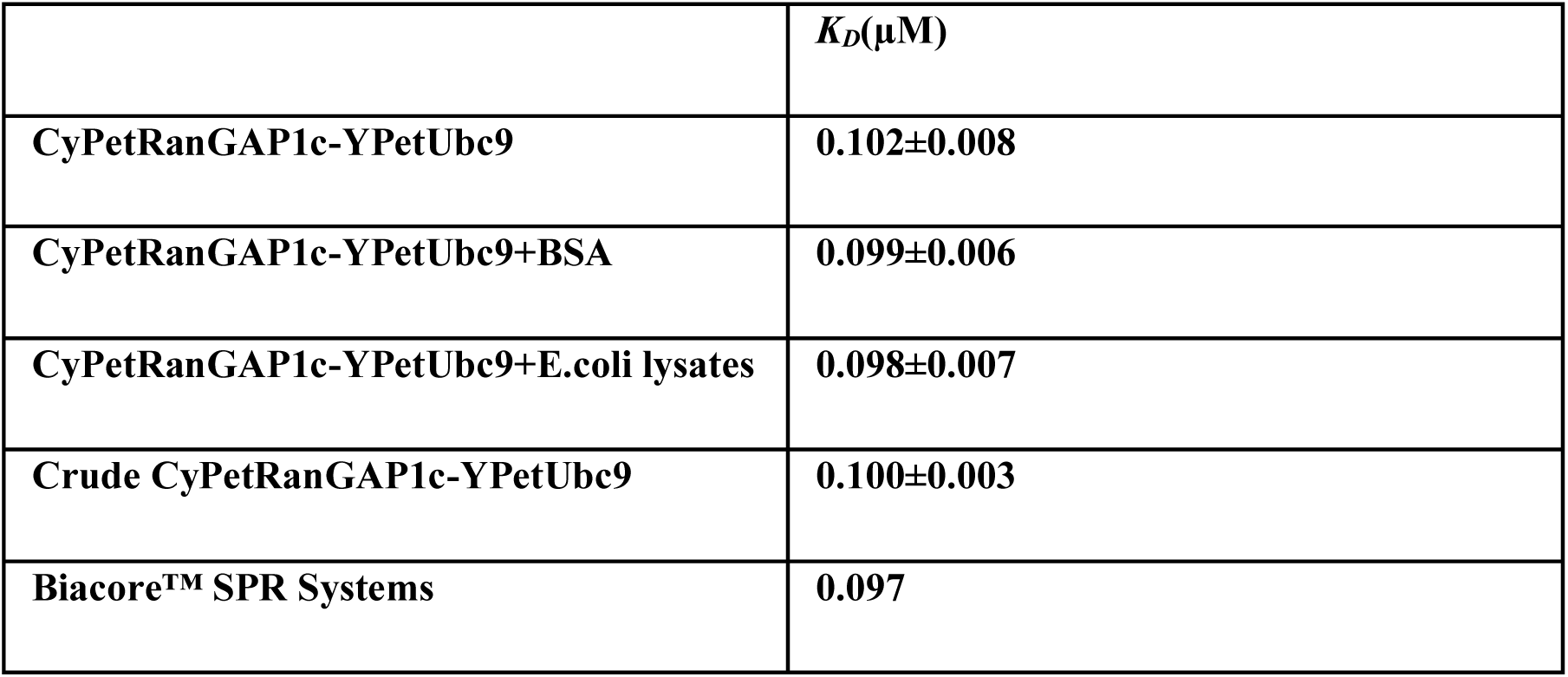
Summary of *K_D_* values of CyPet-Gap1c and YPet-Ubc9 from various conditions and as control, Surface Resonance Plasmon.

### Protein interaction affinity *K_D_* determinations in the presence of other protein(s) and molecules

To test our hypothesis that the **HPAMF** is capable of determining *K_D_* in the presence of the other proteins and even total cell extract, we first determined the *K_D_* value in the presence of BSA and then bacterial cell extract. The CyPetRanGAP1c and YPetUbc9 were purified from bacterial cells using nickel agarose affinity chromatography. We first conducted the assay in four conditions of FRET donor CyPetRanGAP1c, 0.05 µM, 0.1 µM, 0.5 µM, and 1.0 µM, to make sure the reproducibility, and the concentration of YPetUbc9 was changed from 0 to 4 µM with the addition of 1ug of BSA. The Em_FRET_ was determined in the presence of BSA (Fig.2A). The purified proteins and added BSA, 0.1 μM of CyPetRanGAP1c (lane 1, 2, 3), 0.5 μM of CyPetRanGAP1c (lane 4, 5, 6) or 1.0 μM of CyPetRanGAP1c (lane 7, 8, 9) + 1 μM YPetUbc9 (lane 1, 4, 7), 1 μM YPetUbc9 with 1 μg BSA (lane 2, 5, 8), or 1 μM YPetUbc9 with 3 μg BSA (lane 3, 6, 9) (Fig.2B). The values for Em_FRETmax,_ of the mixture in the presence of 1μg BSA were (1.26±0.04)×10^4^, (2.52±0.08)×10^4^, (12.71±0.38)×10^4^ and (24.97±0.76)×10^4^, for concentrations of 0.05, 0.1, 0.5, 1.0 µM of CyPetRanGAP1c, respectively(Supplement Table 1). By plotting Eq.(2) with Em_FRET_ and [YPetUbc9]_total_ in the Prism 5 program, we determined the *K_D_*s from four concentrations of CyPetRanGAP1c (0.05, 0.1, 0.5, 1.0) as 0.098±0.022, 0.092±0.024, 0.105±0.025, 0.102±0.028 µM, respectively(Supplement Table 1). The average *K_D_* is 0.099±0.006 µM. Compared to the *K_D_* without BSA (0.102±0.008 µM), they are very close and no statistically significant difference (Table 1).

**Figure 2.**
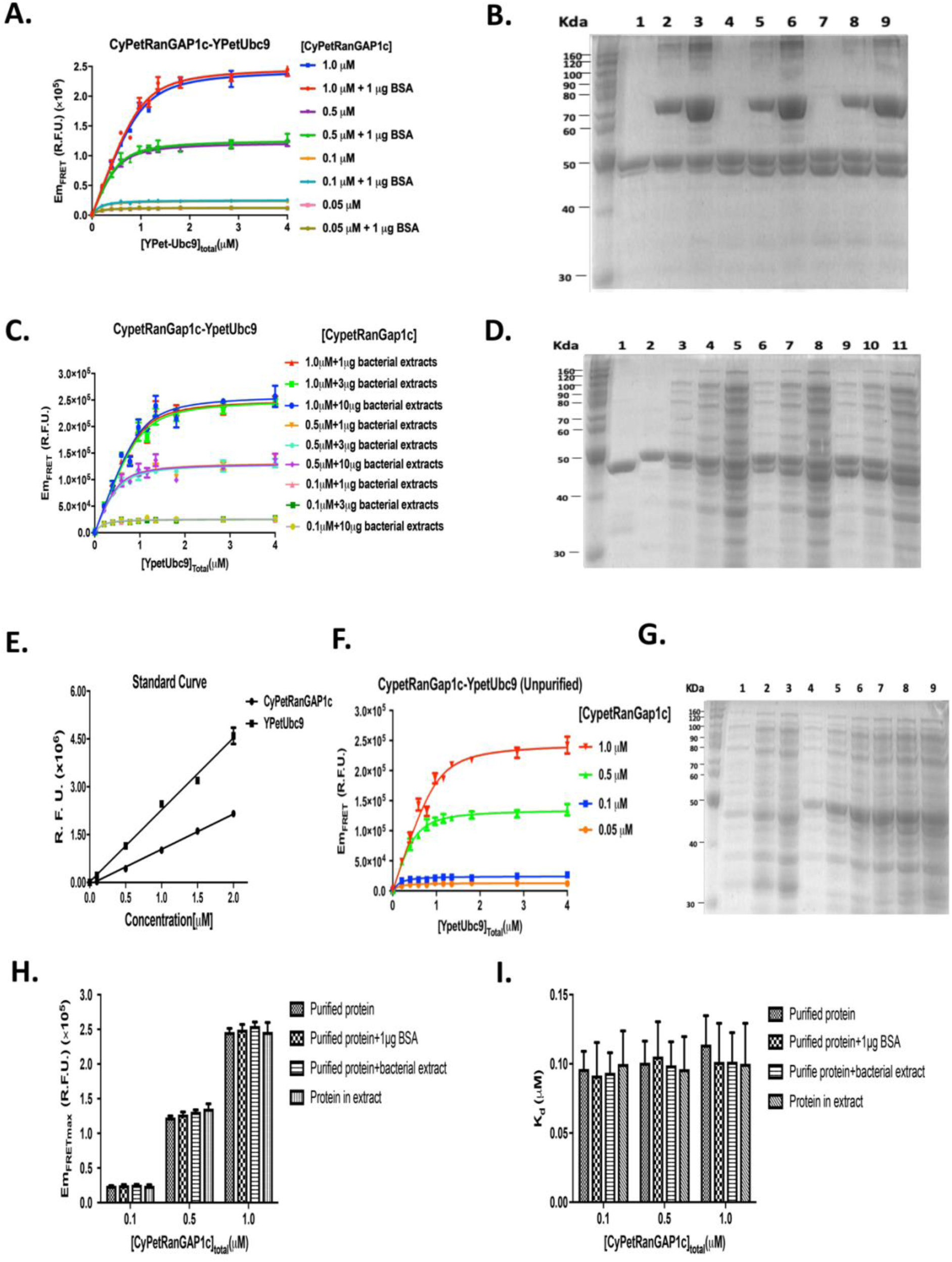
Determine CyPet-RanGAP1c and YPet-Ubc9 interaction affinity *K_D_* in the presence of BSA, or bacterial cell extract or from both extracts. **A.** E_mFRET_ determinations at concentrations of 0.05, 0.1, 0.5, and 1.0 µM of purified CyPetRanGAP1c, respectively, with increasing concentration of purified YPetUbc9 in the presence of 1mg BSA. **B.** The SDS-PAGE protein gel of CyPetRanGAP1c and YPetUbc9 without or with BSA stained with Coomassie. 0.1 μM (lane 1, 2, 3), 0.5 μM (lane 4, 5, 6) or 1.0 μM CyPetRanGAP1c (lane 7, 8, 9) + 1 μM YPetUbc9 (lane 1, 4, 7), with 1 μg BSA (lane 2, 5, 8), or with 3 μg BSA (lane 3, 6, 9). **C.** E_mFRET_ determinations at concentrations of 0.1, 0.5 and 1.0 µM of purified CyPetRanGAP1c, respectively, with increasing concentration of purified YpetUbc9 in the presence of E.coli lysates. **D.** The SDS-PAGE protein gel of CyPetRanGAP1c and YPetUbc9 without or with E.coli cell extract stained with Coomassie. CyPetRanGAP1c (lane 1); YPetUbc9 (Lane 2); 0.1 μM (lane 3,4 and 5), 0.5 μM (lane 6,7 and 8), 1.0 μM (lane 9,10 and 11) of CyPetRanGAP1c + 1 μM YPetUbc9 with 1 μg (lane 3,6 and 9), 3 μg (lane 4,7 and 10), 10 μg (lane 5,8 and 11) of E.coli cell extract. **E.** Standard curve of purified CyPet-RanGap1c or YPet-Ubc9 protein concentrations with its fluorescence emission signal. **F.** SDS-PAGE gel of E.coli cell crude extracts from cells expressing CyPert-RanGap1c (0.1mM, 0.5mM and 1mM in Lane 1-3, respectively), Ypet-Ubc9 (0.1mM, 0.5mM and 1mM in Lane 4-6, respectively), and mixture of CyPert-RanGap1c and Ypet-Ubc9 (0.1mM, 0.5mM and 1mM in Lane 7-9, respectively). **G.** E_mFRET_ determinations of CyPet-RanGAP1c at concentrations of 0.05, 0.1, 0.5 and 1.0 µM with increasing concentrations of YPet-Ubc9 from bacterial extracts. **H.** Maximum E_mFRET_ values of three concentrations of CyPetRanGAP1c with YPetUbc9 from purified proteins, or in the presence of BSA or bacterial extract, or both from bacterial cell extracts. I. *K_D_* values from purified proteins, or in the presence of BSA or bacterial extract, or both from bacterial cell extracts.

The disassociation constants *K_D_* of CyPetRanGAP1c and YPetUbc9 in the presence of bacterial extract were then determined in three concentrations of CyPetRanGAP1c (Fig.2C). From the gel in Fig. 2D, lane 1, 2 are pure CyPetRanGAP1c and YPetUbc9, respectively, 3∼11 are pure CyPetRanGAP1c and YPetUbc9 mixture with 1 µg, 3 µg or 10 µg bacterial lysates proteins, respectively (Fig.2 D). The values for Em_FRETmax_ of the mixture in the presence of E.coli lysates were 2.58±0.074, 13.26±0.43, and 25.21±0.90 when 1 µg E.coli lysates were added, respectively; 2.57±0.079, 13.02±0.39, and 25.19±0.99 when 3 µg E.coli cell lysates were added, respectively; 2.64±0.106, 13.07±0.58, and 26.06±1.14 when 10 µg E.coli lysates were added, respectively (Supplement Table 1).

In all three concentrations of the FRET donor, the protein interaction affinities were determined. When the concentration of CyPetRanGAP1c was fixed as 0.1 µM and 1, 3, 10 µg of the bacterial extract were added to the mixture, the values of *K_D_* were 0.092±0.022, 0.092±0.023, 0.096±0.031 µM, respectively (Supplement Table 1). When the concentration of CyPetRanGAP1c was fixed as 0.5 µM and 1, 3, 10 µg bacterial extract was added to the mixture, the values of *K_D_* were 0.100±0.027, 0.108±0.025, 0.090±0.035 µM, respectively (Supplement Table 1). When the concentration of CyPetRanGAP1c was fixed as 1.0 µM and 1, 3, 10 µg bacterial extract was added to the mixture, the *K_D_*s were 0.093±0.031, 0.109±0.037, 0.103±0.040 µM, respectively. Under these three conditions, the values of *K_D_* were very stable (Supplement Table 1). These results suggest that we can determine *K_D_* using the quantitative FRET assay, and *K_D_* values are very consistent in the absence or presence of BSA or bacterial extract.

So far, we have tested three conditions: one with no contaminating protein, one with single contaminating protein, and another with multiple contaminating proteins. All these conditions gave us similar *K_D_* and Em_FRETmax_ values (Table 1). It indicates that our qFRET-based method can be used to determine *K_D_* under complicated environments, such as in the presence of other proteins. These values of *K_D_* value were directly from two cell extracts without any purification. To achieve this, we first need to know the concentrations of the two FRET pair proteins in the cell extracts. The real concentrations of CyPetRanGAP1c and YPetUbc9 in the crude extracts were determined by comparing their fluorescence signals at 475 nm when excited at 441nm, and 530 nm when excited at 475nm, respectively, with the standard curves of purified CyPet and YPet proteins, respectively (Fig. 2E). The SDS-PAGE gel of the two bacterial cell extracts containing CyPetRanGAP1c and YPetUbc9 proteins, respectively, shown various proteins (Fig. 2F). YPetUbc9 was relatively easier to differentiate because its expression level is higher, while CyPetRanGAP1c is difficult to visualize the band in the gel. The concentrations of CyPetRanGAP1c were fixed at 0.05, 0.1, 0.5, and 1.0 µM, respectively, and the concentrations of YPetUbc9 were increased from 0 to 4 µM. The Em_FRETmax_ was still linear as (1.308±0.041)x10^4^, (2.447±0.075)x10^4^, (13.57±0.39)x10^4^ and (24.63±0.79)x10^4^ (Supplement Table 1). Both Em_FRET max_ and *K_D_*s are very similar to the results from pure protein interaction. The *K_D_*s were 0.102±0.024, 0.100±0.024, 0.096±0.023 and 0.100±0.029 µM, respectively. This result is surprisingly stable and the average of *K_D_* value, 0.100±0.003, agrees with the result from the pure proteins (Table 1).

Further analysis shows no statistically significant difference among these Em_FRETmax_ and *K_D_* values in all the four conditions, demonstrating the reliability and robustness of the FRET-based *K_D_* determination approach using un-purified proteins (Fig. 2H and I). This demonstrate the HPAMF method is very reliable and robustness.

### HPAMF shows a high affinity of the SUMOylation E3 ligase with the E2 conjugating enzyme, conferring a specific pairing of the E3-E2 in the cascade

To test the ability of HPAMF in a very challenging situation, such as the interaction affinity of SUMOylation E3 ligase and E2 conjugating enzyme, we applied the HPAMF to determine the affinity of PIAS1 and E2 Ubc9. *In vitro*, with the help of the E1 enzyme, the E2 enzyme Ubc9 can recognize and transfer SUMO peptides to most substrates directly. In contrast, the E3 ligases, such as PIAS1, are speculated to facilitate most SUMOylation processes by gluing the E2∼SUMO and substrate together to promote SUMO peptide transfer *in vivo*. Although genetics and non-quantitative biochemical experiments have shown that PIAS1 is an E3 ligase for SUMOylation, it has never been shown from a kinetics perspective that PIAS1 interacts with Ubc9 at high affinity, mostly due to the difficulty of obtaining a large quantity of E3 PIAS1 protein in many expression systems.

We, therefore, applied the HPAMF to determine the affinity of PIAS1 and Ubc9 interaction. We cloned the PIAS1 cDNA after codon optimization and Ubiquitin E2 enzyme the Ubc12 cDNA as control, as fusion proteins with CyPet and YPet, respectively, into pET28(b) vector. We optimized the codons of PIAS1 because almost no PIAS1 protein could be expressed and purified with its original codons. The plasmids were transfected into E.coli Bl_21_(DE3) cells and induced overnight with IPTG. The protein expressions were examined with SDS-PAGE gel for samples of uninduced and induced cells and supernatants after centrifugation of sonicated cells (Fig, 3A). The CyPet-Ubc9 and CyPet-Ubc12 were well induced (Figure 4A, lane1-3 and 7-9), while the YPet-PIAS1 protein was hardly seen (Figure 4, lane 4-6). We then calculated the CyPet-Ubc9 and CyPet-Ubc12, and YPet-PIAS1 amounts using external CyPet and YPet standard curves, respectively. We then performed the FRET titration experiments. In each titration experiment, the concentration of CyPet-Ubc9 or Cypet-

Ubc12 was fixed and increasing concentrations of YPet-PIAS1 were added. The values of Em_FRERT_ in each point were calculated according to Eq.(2). The titration curves of Em_FRERT_ are shown (Fig. 3B). The titration curves of CyPet-Ubc9 and YPet-PIAS1 shown good dose-dependent increase with the concentrations of CyPet-Ubc9, while the titration curves of CyPet-Ubc12 with YPet-PIAS1 did not show any Em_FRERT_ signal even at the highest concentration of YPet-PIAS1, suggesting a strong specific interaction between Ubc9 and PIAS1. We then determined the interaction affinities of CyPet-Ubc9 with YPet-PIAS1 at each concentration of CyPet-Ubc9 (Figure 4C). The *K_D_* values were very consistent, ranging from 0.26 ± 0.04 to 0.30 ± 0.04 from the four testing conditions. This result suggests that PIAS1 interacts with Ubc9 at very high affinity and, therefore, sets up the basis of specificity. Further analysis shows no statistically significant difference in these *K_D_*values, again demonstrating the reliability of the HPAMF.

**Figure 3.**
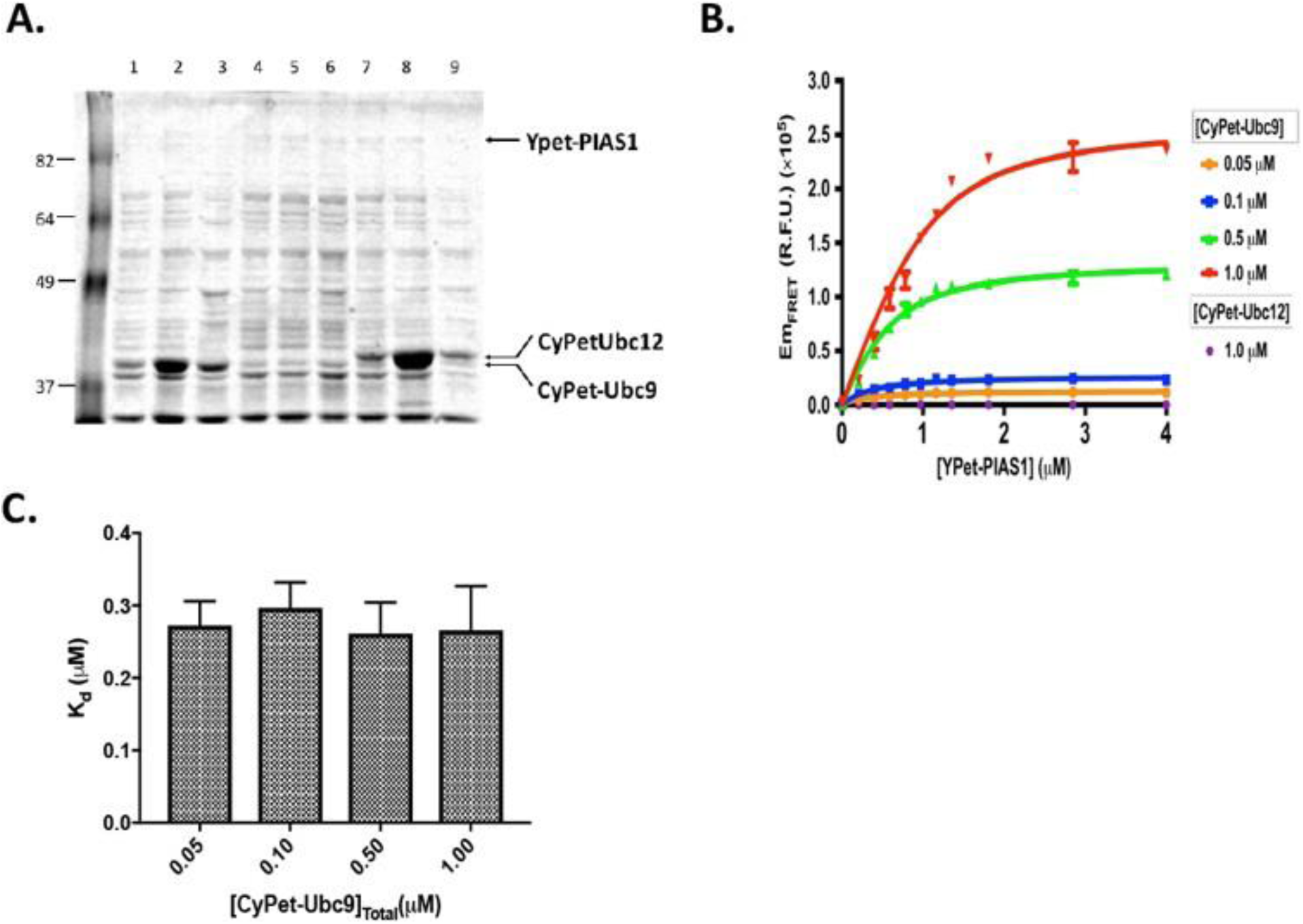
Determine specificity of SUMO E3 ligase, PIAS1, to its E2 conjugating enzyme, Ubc9, from interaction affinity. **A.** SDS-PAGE gel of E.coli cells of un-induced (lane 1, 4,and 7), induced (lane 2,5,and 8) or supernant (lane 3,6, and 9) of CyPet-Ubc9, YPet-PIAS1, CyPet-Ubc12, respectively. **B.** Em_FRET_ determinations of various concentrations of CyPet-Ubc9 or CyPet-Ubc12 with increasing concentrations of YPet-PIAS1. **C.** *K_D_* determinations of PIAS1 with Ubc9 from various concentrations of CyPet-Ubc9.

**Figure 4.**
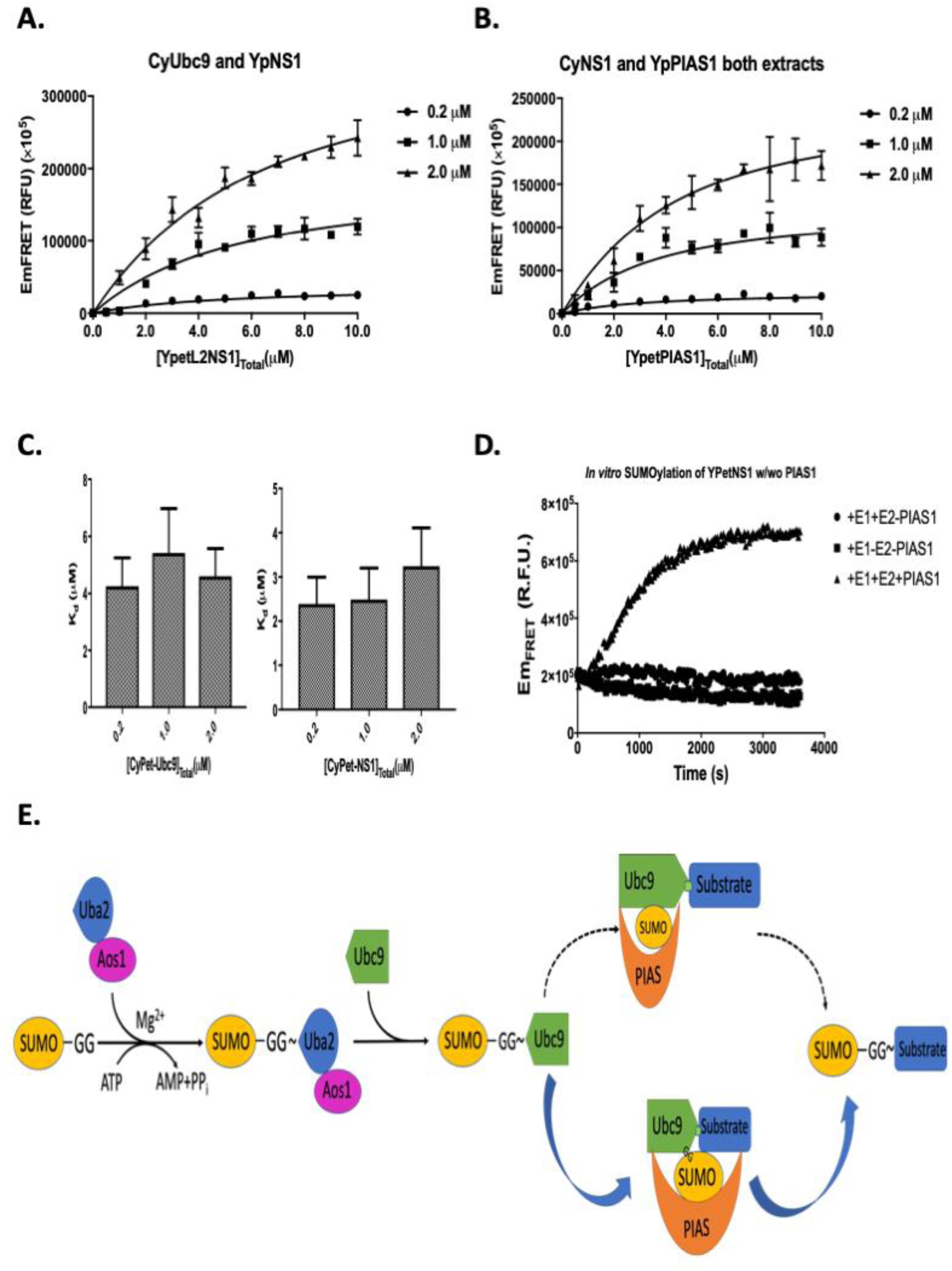
Both SUMO E2 conjugating enzyme E2 and E3 ligase contribute to substrate recognition. **A.** E2 Ubc9 recognizes influenza virus protein NS1 at high affinity. **B.** E3 PIAS1 recognizes influenza virus protein NS1 at high affinity. **C.** The bar graph of *K_D_* values of NS1 with Ubc9 and PIAS1, respectively. **D.** PIAS1 facilitates the SUMOylation of NS1. E. The model of substrate recognition in the SUMOylation cascade.

### SUMO E3 ligase recognizes influenza virus protein NS1 directly with high affinity

It has been suggested that there may be two possible models on how the SUMOylation cascade can transfer the SUMO peptide to substrates: the first one was that both E2 and E3 would interact with substrates simultaneously; the second was that only E2 would interact with substrates, while E3 only brings E2 and SUMO peptide together to the substrates, and E3 does not recognize substrates. To distinguish these two models from the kinetics perspective, the HPAMF method was used to determine the interaction affinities of SUMOylation E2 Ubc9 and E3 PIAS1 to influenza virus protein NS1. The CyPet-Ubc9 and YPet-NS1 proteins were obtained from bacterial Bl_21_(DE_3_) cells. The Em_FRET_ of CyPet-Ubc9 and YPet-NS1 was determined in three concentrations of CyPet-Ubc9 (Fig. 4A). Then, we derived the interaction affinity of PIAS1 with NS1. The YPet-PIAS1 and CyPet-NS1 were expressed in bacterial cells, and supernatant mixtures were prepared after sonication. According to the CyPet and YPet standard curves, the concentrations of YPet-PIAS1 and CyPet-NS1 were determined. Then we determined the Em_FRET_ in three concentrations of CyPet-NS1, 0.2μM, 1.0μM and 2.0μM, respectively (Fig. 4B). The average *K_D_* value was 2.96 ± 0.47 uM, slightly higher than that of UBC9 to NS1, which average was 4.84 ± 0.64 μM (Figure 5C). This result suggests that, although both Ubc9 and PIAS1 interact with the substrate at modest affinities, they collaborate to contribute to substrate recognition and confer higher affinity and specificity.

To determine the contribution of SUMO E3 ligase PIAS1 during the SUMOylation process, we then carried out a full SUMOylation assay with or without PIAS1 using FRET assay. In this setting, if the SUMOylation happened, the conjugation of CyPet-SUMO1 to YPet-NS1 from the SUMOylation cascade would lead to FRET signal generation. The SUMOylation assay was carried out with CyPet-SUMO1, E1(Aos1/Uba2), E2(Ubc9), and YPet-NS1 in the presence or absence of the E3 ligase, PIAS1. In this assay with a low concentration of Ubc9(1uM), the SUMO conjugation only happened in the presence of PIAS1, indicating a critical role of E3 ligase in the SUMOylation reaction at a low protein concentration environment. As a control, no FRET signal was observed without ATP (Fig. 4D). This result indicates that PIAS1 can help the SUMO conjugation and may reflect physiological conditions in which PIAS1 has been discovered to be required for some substrate SUMOylation *in vivo*(*37*).

From these data, we propose the following mechanistic model of the SUMOylation process (Fig.4E). After the activation of SUMO peptides by E1 and E2 enzymes, the SUMO peptides are conjugated to substrates with the substrate recognitions by both E2 and E3 enzymes, therefore, conferring high affinity and specificity. This model, supported by kinetics parameters, may explain the long-term mystery of SUMOylation specificity *in vivo*.

## DISCUSSION

In these studies, we present a novel approach (HPAMF) to determine protein interaction affinity directly from cell extracts without any purification using qFRET analysis and discover a kinetics basis for the mechanistic understanding of SUMOylation substrate recognition from E2/E3 co-recognition mechanism. We systematically developed and validated the HPAMF technology for protein interaction affinity determination for purified proteins in the presence of BSA and cell extract and interactive proteins in cell extracts without any purification. This HPAMP method can accurately determine protein interaction affinity in an environment with other proteins and even without purification in a high-throughput mode, representing a major methodology advancement and the capability of determining protein interactions quantitatively on a large scale.

The determinations of interaction affinities between SUMOylation E2, Ubc9, and E3, PIAS1, enzymes with the substrate, influenza NS1 protein, could provide an explanatory mechanism underlining the nature of SUMOylation substrate recognition to achieve high substrate specificity *in vivo*. There are approximately 6000 SUMO substrates but only one SUMO E2 conjugating enzyme in the human genome(*38*). How the SUMOylation achieves high specificity remains a major enigmatic issue. If substrate recognition solely depends on E2-substrate interactions, achieving a high specificity of SUMOylation, which is essential for many physiological processes, is very challenging(*31*). If the SUMOylation E3 ligase can also provide a high interaction affinity with substrates, E3 ligases alone or with E2 conjugating enzyme can bring the substrates to the SUMO-E2 complex for SUMO transfer in a high specificity. However, the interaction affinity of SUMOylation E3 ligase with substrates was not determined before, probably due to challenges in E3 ligase expression and purification, as evidenced by the fact that only a limited SUMOylation E3 ligases have been crystallized so far(*31, 32, 39*). The two-enzyme recognition model of SUMOylation proposed in this study leads to a significant step toward understanding SUMOylation substrate recognition *in vivo*.

Among the methods evaluated, our HPAMF approach is the only one capable of determining protein interaction affinity without purification and in high-throughput mode. The HPAMF also provides kinetics parameters of multi-protein reactions in protein cascade reactions. The additional ability to link protein interaction affinity with multi-protein reaction kinetics provides an opportunity to investigate biochemical reactions in a complex environment. Extension of qFRET approaches to biochemical reactions would allow generations of far more comprehensive quantitative maps of biochemical reactions in life. Further applications of the HPAMF to diverse protein-protein interactions, especially those in which expression and purification have been limited, should help understand vast protein interactions that are currently unknown and provide a universal measurement for quantitative systems biology.

Many methodologies and technologies have been developed to determine protein-protein interaction affinity due to its fundamental role in all physiology processes and pharmaceutical developments. In this study, we also verified our results using the SPR method, demonstrating that the novel qFRET assay produces consistent results that agree well with those determined using classical *K_D_* measurement technologies. The SPR itself may have some potential intrinsic drawbacks. For example, purified proteins may lose their native conformation and heterogenicity after immobilization on the sensor surface, or fixed binding orientation may block the interactive surface and decrease affinity(*40, 41*). In contrast, the HPAMF method has already considered the potential orientation problem as it determines protein interactions in solution. Also, due to the surface immobilization of the ligand, the local concentration is much higher than it is in the solution, and the mass transfer effect may induce binding kinetic distortion from ideal pseudo-first-order binding(*41, 42*). In addition, nonspecific binding to the sensor chip and rebinding effect may occur, and thus, interfere with the accuracy of *K_D_* value(*43*). Furthermore, the SPR method for *K_D_* determination might not be reliable if a protein-protein interaction does not follow the Langmuir-type binding model(*42, 44*). In addition, the SPR assay needs 40 min also to get the final result. So, it won’t be feasible for large sample tests or high-throughput assay. If the final product formation result in interaction, it may produce inconsistent results as enzymes may lose activities during this period of time. In contrast, six concentrations in triplicate only takes less than 1 minute in the HPAMF method. In addition to SPR, many different methods, including isothermal titration calorimetry (ITC), ultracentrifugation, and radioactive labeling, have also been used for *K_D_* determination(*3, 4*). These methods offer standard experimental procedures but also hold some disadvantages. They often require expensive instrumentations, environmentally unfriendly labeling, and especially can’t determine impure protein interactions. Isothermal titration calorimetry (ITC) requires relatively large amounts (i.e., minimolar range) of proteins, and systematics errors in heart calibration, cell volume and other issues, such as baseline errors and gas bubbles, can lead to inaccurate results(*45–47*). The ITC also needs relatively expensive specialized equipment. In ultracentrifugation assay, the long centrifugation time can perturb the equilibrium between bound and free proteins due to centrifuge force, especially if the protein interaction dissociation rates are fast. Thus, the determined *K_D_* values do not represent true equilibrium constants. In addition, peripheral or membrane proteins can bind nonspecifically to the test tube wall during centrifugation, which leads to fewer free proteins in binding events. The fluorescent polarization (FP) approach can also determine the protein interaction dissociation constant. In principle, the FP approach can potentially address the above issues. However, the FP approach also needs purified interacting partners and the determined *K_D_* is inaccurate if the fluorescence-labelled partner is too big as it measures the polarized fluorescent signal(*48*).

The consistent results of the *K_D_* determination at the micromolar range from un-purified proteins show that the HPAMF method is not only sensitive at the affinity of nanomolar level but also accurate and reliable using a small amount of the interactive proteins. This may provide a very beneficial advantage in practice, especially for some precious proteins or proteins in the R&D process without a large amount of protein available, such as antibody drug discovery. Furthermore, other approaches for *K_D_* determination, such as SPR or ITC, need multiple steps and a relatively large amount of purified proteins and are hardly implemented as a high-throughput assay due to limitations in both instrument capability and available protein. While the FRET assay has become more prevalent in basic research, diagnosis, and drug discovery, the HPAMF method would advance FRET technology to a more robust technology for quantitative and high-throughput biological/biomedical research and pharmaceutical development.

## ACKNOWLEDGEMENTS

We are very grateful to Dr. Michael Pirrung in the Department of Chemistry, University of California at Riverside for allowing us to access his molecular biology lab. We thank all the members in Liao’s group for their very close collaborative work and assistance with this study.

## AUTHOR CONTRIBUTIONS

L,J conduct the initial experiments and wrote the original manuscript; Z. X. repeated and validated the experiments; Y.L. conducted the qFRET analysis; J.Li provided statistic advices; J.Liao. came the original idea and supervised the studies, and wrote the manuscript.

---

**Supplement Table 1.** Summary of the maximal FRET emission Em_FRETmax_ and *K_D_* values in different conditions.

| CyPetRanGAP1c(μM) | $K_D$(μM) | $Em_{FRETmax}$ (R.F.U.)($\times 10^4$) |
| --- | --- | --- |
| 0.05 | 0.098±0.014 | 1.227±0.022 |
| 0.1 | 0.096±0.013 | 2.433±0.041 |
| 0.5 | 0.101±0.016 | 12.29±0.23 |
| 1 | 0.114±0.021 | 24.61±0.53 |
| 0.05+1 μgBSA | 0.098±0.022 | 1.260±0.036 |
| 0.1+1μg BSA | 0.092±0.024 | 2.523±0.080 |
| 0.5+1 μg BSA | 0.105±0.025 | 12.71±0.38 |
| 1.0+1 μg BSA | 0.102±0.028 | 24.97±0.76 |
| 0.1+1μg bacterial extract | 0.092±0.022 | 2.583±0.074 |
| 0.1+3μg bacterial extract | 0.092±0.023 | 2.572±0.079 |
| 0.1+10μg bacterial extract | 0.096±0.031 | 2.636±0.106 |
| 0.5+1μg bacterial extract | 0.100±0.027 | 13.26±0.43 |
| 0.5+3μg bacterial extract | 0.108±0.025 | 13.02±0.39 |
| 0.5+10μg bacterial extract | 0.090±0.035 | 13.07±0.58 |
| 1.0+1μg bacterial extract | 0.093±0.031 | 25.21±0.90 |
| 1.0+3μg bacterial extract | 0.109±0.037 | 25.19±0.99 |
| 1.0+10μg bacterial extract | 0.103±0.040 | 26.06±1.14 |
| 0.05 (from crude extract) | 0.102±0.024 | 1.308±0.041 |
| 0.1 (from crude extract) | 0.100±0.024 | 2.447±0.075 |
| 0.5 (from crude extract) | 0.096±0.023 | 13.57±0.39 |
| 1.0 (from crude extract) | 0.100±0.029 | 24.63±0.79 |
| SPR Method *by Ling* | 0.097 |  |

**Supplement Figure 1.**
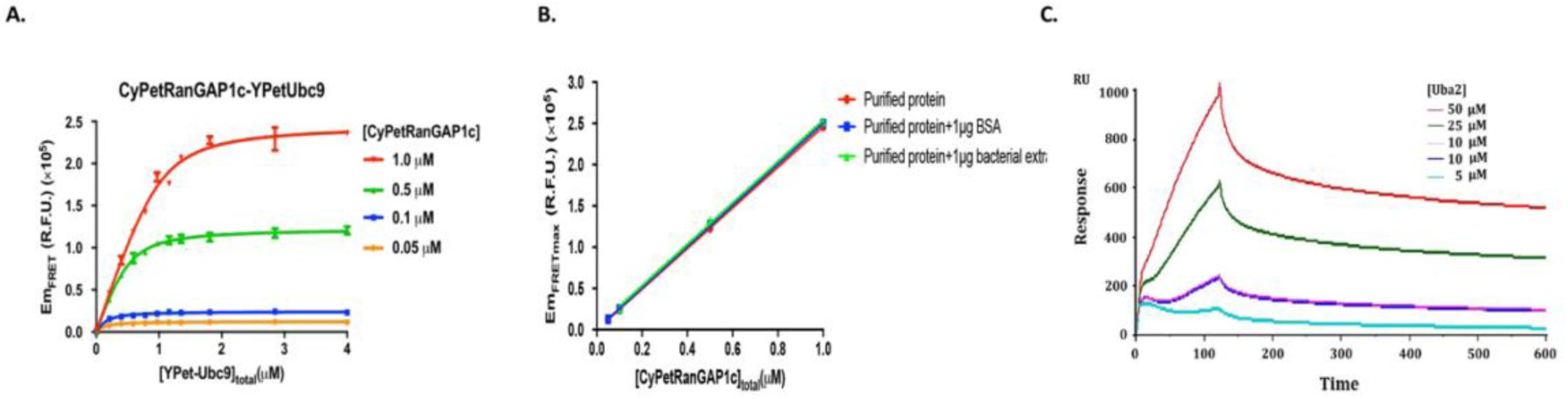
Interaction affinity determination of CyPet-Ubc9 and Ypet-PIAS1. **A.** Em_FRET_ determinations of purified CyPet-Ubc9 at concentrations of 0.05mM, 0.1mM, 0.5mM and 1.0mM with increasing concentrations of purified Ypet-PIAS1. B. Determinations of Em_FRETmax_ at different concentrations of the donor CyPet-RanGAP1C with YPet-Ubc9 in the absence and presence of other proteins. The maximal FRET emission is proportional to the amount of CyPet-RanGAP1 in the assay with or without other proteins. C. Dendrogram CyPet–RanGAP1c and YPet–Ubc9 interaction by the surface plasma resonance (SPR).

